# Software for agriculture climate risk management focused in smallholders

**DOI:** 10.1101/745109

**Authors:** Priscila Pereira Coltri, Hilton Silveira Pinto, Yasmin Onorio de Medeiros, Kaio Shinji Hashimoto, Giovanni Chaves Di Blasio, Eduardo Lauriano Alfonsi, Renata Ribeiro do Valle Gonçalves, Waldenilza Monteiro Alfonsi

**Author notes:** Cidade Universitaria “Zeferino Vaz”, Campinas/SP, Brazil. 13083-970.

## Abstract

**T**aking Persimmon (*Diospyros kaki* L.), Fig (*Ficus carica* L.) and Papaya (*Carica Papaya* L) fruits in São Paulo State, southeast of Brazil, as a case of study, we present here new software that was designed to support smallholders to manage their climate risk on the production area. The main idea of this new software named “Brazilian Mapping for Agricultural Zoning System” (BRAMAZOS) is to transform scientific knowledge into useful information for crop climate risk management, indicating the risk of crop failure and which is the limiting meteorological element for the area unsuitability. The software was developed based on user experience design, focusing on the users facilities with a friendly interface. We simulated these fruits climate risk in current and future scenarios of climate change using ETAHadgem ES Regional Climate Models (RCM), which is a downscaling of the Global Climate Model HadGEM2-ES, based on the IPCC 5th Assessment Report (AR5). We observed changes in the climate risk zone distribution for Persimmon and Papaya cultivation, which could lose almost 30% and 9% of the suitable area respectively. On the other hand, fig did not present significant reduction in the climate risk zone. The results presented here suggested that the temperate fruit examples used here seem to be more sensible to the temperature increase and, the tropical example seems to be more precipitation decrease sensitive. We discussed the significance of BRAMAZOS software as tool to support efficient information to climate risk management, providing agroclimatic information that are efficient to assist decision making, increase food security with the intention to reduce the climate impact on smallholders development and resources management issues.

## 1. Introduction

According to Food and Agriculture Organization of the United Nations (FAO), Brazil is globally recognized as one of the major agricultural powerhouse and accounts for one fifth of the global food production. In Brazil, agribusiness system is responsible for about one third of the country economy and a crucial contributing sector for economic growth and foreign exchange. Due to the country size, the agricultural sector is heterogeneous in terms of climate, geographic conditions, infrastructure and economic investments across the regions. Brazilians farmers face a number of risks to their production process as price volatility, economy, marketing, public politics, among others. Most of the national agriculture is constrained by climate conditions, making events such as drought and excessive rainfall a major source of production risks (Cunha et al., 2001).

To address these challenges, minimizing Brazilian agriculture losses, in 1996, the National Government published the Official Agricultural Climate Risk Zoning, an instrument of agricultural economic policy that was designed to minimize risks related to adverse climatic phenomena. This program allows each municipality to identify the best plant season of the crops, in three different types of soil and crop cycle (Cunha et al. 2001). After this program implementation, the national grain production increased and the losses caused by climatic advertise reduce significantly. According to the Brazilian Ministry of Agriculture, in 1991 the grain production in Brazil was about 57.9 million tons, cultivated in an area of 37.9 million ha, and, in 2010, the production increased up to almost 150 million tons in an area of 47.5 million ha. The increase in production was about 153.7 % (4.8% by year) while the total cultivated area increased only 25.4% (1.7% by year). The evolution of technology was the responsible for this gain in productivity and the Agricultural Climatic Risk Zoning, after becoming a public policy in the country, certainly contributed for this fact.

Public programs similarly the Agricultural Climate Risk Zoning is efficient because agriculture is an economic activity completely connected with weather and climate conditions. Precipitation and temperature can explain approximately 70% of the variation of the total factor productivity (Liang et al. 2017). Deciding what to plant, when to plant and where to plant, with lower probability of crop failure risk, is a direct function of weather persistency in the phenological plant cycle. Cultivated plants have a suitable climatic range in which they grow and develop economically; therefore, the previous knowledge of the climatic characteristics of the cultivated areas is necessary for the crop success or failure.

The relationship between climatic elements and the agriculture production is not trivial (Camargo, 2010) but some agrometeorological information as the products from water balance (soil water available, evapotranspiration, net water requirements) associated to whether conditions (air temperature, precipitation pattern, frost probability, drought frequency, radiation, wind speedy) contributes to calculate climate risk in agriculture.

Climate Risk Management (CRM), in agriculture, is a process that advise decision making through the application of climate knowledge, in a multidisciplinary scope, with the intention to reduce the climate impact on agriculture development and resources management issues (Hellmuth et al., 2009). This approach is necessary to indicate land suitability, minimizing ecological damages, improve decision making and cooperating to sustainable agriculture. This method increases the knowledge on a full range of crops to be cultivated in different regions, especially in the possible climate change scenarios, which is one of the most important challenges to food production and security nowadays.

According to IPCC, climate system is changing. In 2014, last scientific assessment report (AR5) announced that the global warming is unequivocal, changing the climate system, increasing temperatures and changing precipitation pattern, intensifying extreme events and, consequently, modifying the water balance of innumerous producer regions. Therefore, new challenges appear to agriculture. Since the first assessment report (AR1) from IPCC, in 1990, the scientific research focusing on the impact of the climate change on agriculture has grown deeply and significantly. Particularly in Brazil, the last two decades have seen important efforts to understand this possible changes in local agriculture and, researches have been announcing the impact of the possible climate change in productivity areas, losing suitable areas (Assad et al., 2004; Zullo Junior et al., 2006; Pinto et al., 2008), production (Tavares et al., 2018), increasing pest and disease vulnerability (Ghini et. al., 2008; Ghini et al., 2011) and reporting the importance of adaptation and mitigation activities (Coltri et al., 2015). Most of these studies were concentrated in principal commodities as coffee, corn, rice, potatoes, wheat, sugarcane and soybean. Few studies focused on fruit production, which is an important source of income of the family farmers and smallholder in the world, and where these impacts could be particularly severe because of the high vulnerability of these farmers (Donatti et al., 2018; Holland et al., 2017; Harvey et al., 2018)

Smallholders, or family farmers, play an important role in agriculture system and world food security (FAO, 2014). It is estimated that 85% of the world farm is constituted by smallholders farmers (Harvey et al., 2014). In Brazil, this segment also represents more than 80% of the production units, growing every year (Herrera et al., 2017). Nowadays, Brazil has nearly 4.7 million smallholders who occupying 89 million hectares, being a subsistence source for 17 million people (Herrera et al., 2018). This sector, represented in 2006, 38% of the gross value of agriculture (Herrera et al., 2018) and it is normally related to subsistence agriculture (Morton, 2007). Even with this economic and social importance, family farming is still a vulnerable sector, because it is complex, diverse and risk susceptible (Chambers et al., 1989) and normally characterized by low levels of investment. This segment come is considered the most climate vulnerable group, undermining income security and the total productivity (Abdul-Razak et al., 2017). Smallholders normally have small participation on public programs, especially because there are lacks of information (Souza-Esquerdo et al., 2014) and, often, the knowledge does not reach the family farmer. In the climate change context, it is an urgent need to identify adaptation options to this important sector.

One important approach to reduce smallholders vulnerability is access to knowledge, offering simple and robust tools for their own decision making, relating climate to agronomic impacts and allowing the farmers to manage their local climate risk. Land suitable evaluation knowing the climatic potential of a region for the development of plants and translating agroclimate information in useful information, is especially significant for this end. However, few tools are available to this important sector, allowing the climate risk management. Although the Official Agricultural Climate Risk Zoning is a national program, do not present an adequate scale and detail of the information to manage their particular scenarios and risks.

Therefore, we propose here a new system that was designed to support smallholders to manage their climate risk on the production area. The main idea of this new program is to transform scientific knowledge into useful information for crop climate risk management, indicating the risk of crop failure and, thus, permitting smallholders to be better-informed to initiate agriculture operations. The software called “Brazilian Mapping for Agricultural Zoning System” (BRAMAZOS) was developed based on user experience design, focusing on the users facilities with a friendly interface. In this study, we presented the climate risk analyses to temperature and tropical fruits in São Paulo state, southeast of Brazil, in actual and future climate scenarios, indicating the possible tools to these agricultures manage the climate risk.

## 2. Material and Methods

BRAMAZOS software was developed at Center for Meteorological and Climate Research Applied to Agriculture (CEPAGRI), at University of Campinas, Brazil. The purpose of the software is to give a table or/and a map corresponding to the climatic risk and the climatic suitability for specific fruits at Brazilian states. We developed the system based on SARRA software (Système d’analyse régionale des risques agroclimatiques) (Baron et al. 1999), created by CIRAD (Centre de coopération internationale en recherche agronomique pour le développement). SARRA is package software designed to answer the needs of non-specialists in agroclimatology and information, planning the most appropriate planting data and irrigation.

SARRA is composed by three packages: SARRAMET, SARRABIL and SARRAZON (Baron et al., 1999). SARRAMET aims to study the relationships between climate and plant physiology, SARRABIL is responsible by the water balance and the irrigation scenarios, working on a small scale. SARRAZON is responsible for the analysis of the regional zoning, indicating the agriculture potential and contributing to the best agricultural yield. We used SARRAZON basis to develop BRAMAZOS software.

### 2.1 Climatic Data

Climate data is an essential database to run the BRAMAZOS software. Therefore, we created BRAMAZOS climate database, which contains, nowadays, two categories of climate data: real data (from meteorological station) and climate model data. In this new climate database, there is space to new sources of climate data (eg. different climate models, satellites data, etc).

For the simulation presented here, we used real meteorological data and climate model data. Agritempo database (www.agritempo.gov.br) was used as the real climate data, which contains 86 meteorological station at São Paulo state, with daily temperature (minimum, maximum) and rainfall data. On average, these stations have 10 years of weather data.

To the climate model, we choose the Regional Model EtaHadGEM2-ES (Chou et al., 2014), because the spatial resolution is preferable than the Global Climate Models (GCM) being refined enough to capture local characteristics (Tavares et al., 2018), which is more indicated to study small and local farmers. EtaHadGEM2-ES has 20km of spatial resolution, and is a downscaling of the Global Climate Model HadGEM2-ES, based on the IPCC 5th Assessment Report (AR5).

We used three climatic scenarios from the model:

i. current climate (data from 1961 to 1990);
ii. future climate data based on the Representative Concentration Pathway scenarios RCP 4.5 Wm^−2^ radiative forcing, which correspond to the optimistic greenhouse gases emission scenario, from 2011 to 2040 and,
iii. future climate data based on the Representative Concentration Pathway scenarios RCP 8.5 Wm^−2^ radiative forcing, which correspond to the pessimist greenhouse gases emission scenario, from 2011 to 2040.

The real climate data was used to evaluate current model data. However, Agritempo system presented a higher number of meteorological stations with temperature series failures. Therefore, to compare the current scenario (1961-1990) from EtaHadGEM2-ES to real data, we used regression equations proposed by Pinto et al (1972) (equation 1), developed to São Paulo state, to estimate the annual temperature average. The equation 1 uses altitude and latitude to estimate the temperature.

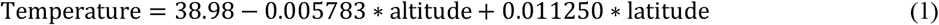

We verified that EtaHadGEM2-ES underestimate the average annual temperature in 2°C to São Paulo State. Therefore, to run BRAMAZO software we add 2°C in current and future scenarios.

To evaluate EtaHadGEM2-ES precipitation, we chose meteorological station that presented less than 5% of rainfall data failures. We used 46 meteorological station and compared to the same coordinate point (latitude and longitude) extracted from the model. Therefore, to run the model, we decreased the total daily precipitation by 20%.

### 2.2 Model Development

#### a) BRAMAZO Models – Structure and Implementation

The BRAMAZO program nowadays, is available via web page and accessed by an internal network at Centre for Meteorological and Climate Research Applied to Agriculture (CEPAGRI). The conceptual flowchart of the software is presented on Figure 1. The theoretical idea is that the user starts the simulation choosing the culture of interest, soil group, Brazilian state and the climatic database (real data, model data, future optimist scenario, future pessimist scenarios, etc) to manage the local climate risk.

**Figure 1.**
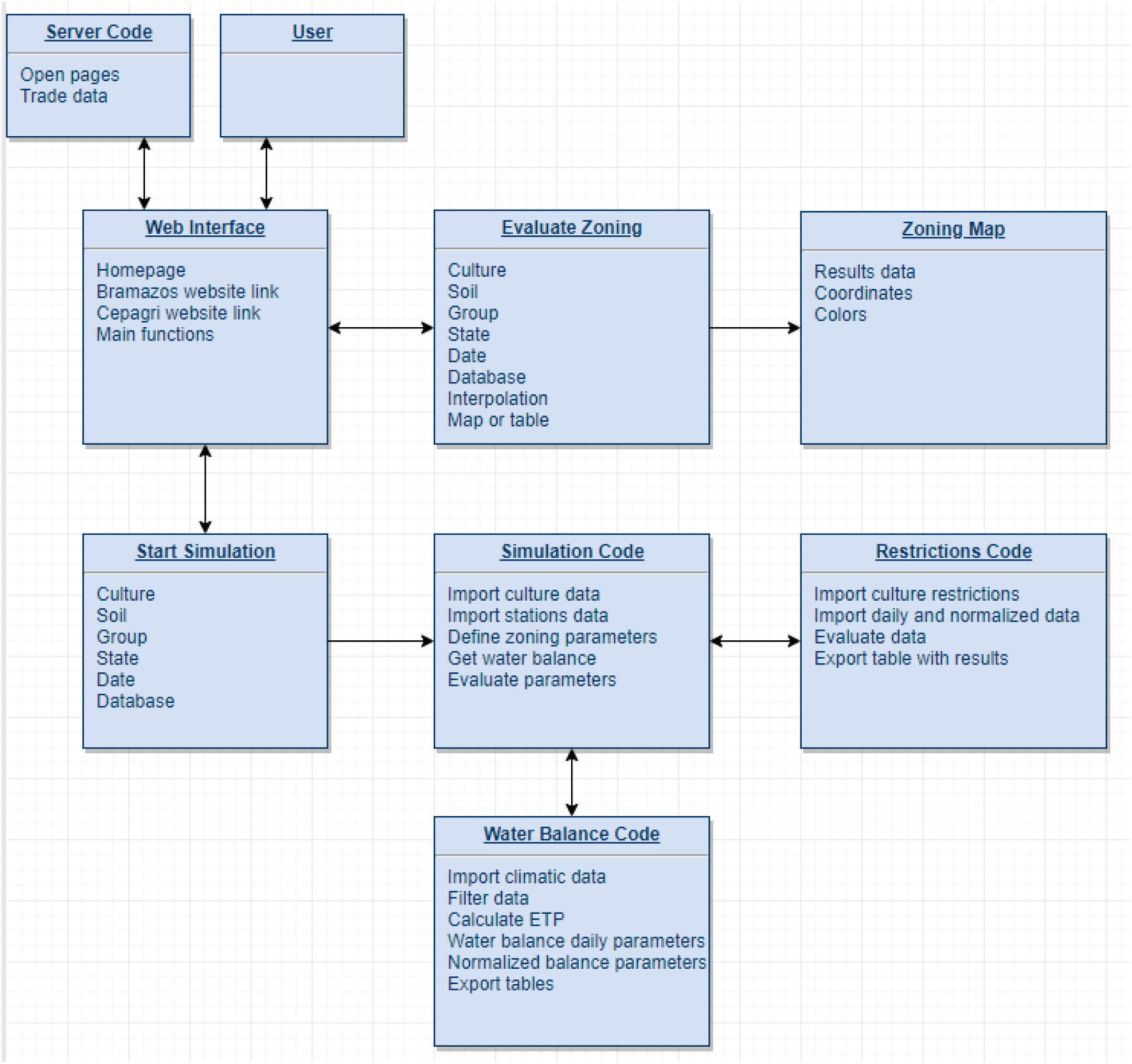
Conceptual flowchart of BRAMAZO software

Until now, the BRAMAZO is able to run the climate risk management to five annual crops (cotton, rice, beans, corn and soybean) and thirteen perennial crops (banana, coffee, pineapple, persimmon, citrus, fig, papaya, passion fruit, nectarine, peach, mango and manioc). The main restrictions implemented are: water requirement satisfaction index (WRSI), average minimum temperature on phenological phase I (Tmin), average mean temperature (Tmed), average maximum temperature on phenological phase III (Tmax), minimum precipitation required (Prec), annual mean temperature (TMA), frost risk, annual water deficiency (DHA), monthly mean temperature (TMM), number of cold hours (NHF), altitude (ALT), annual precipitation (PMA), monthly water deficiency (DHM) and water index (IH).

The software functions are split into two: generate analysis and evaluate data. The first one calculates the zoning parameters per station and the second one spatializes the results interpolation and shows them on a map. The interpolation method applied is the square inverse distance weighting and it takes into consideration latitude, longitude and altitude.

Figure 2 shows the schematic for the program behavior, regarding web pages and python classes connection. Note that the user can access the homepage and from there go to the page for starting simulation or get the zoning map. In addition, there are links for CEPAGRI web page and BRAMAZOS information, such as the team members and a small documentation.

**Figure 2.**
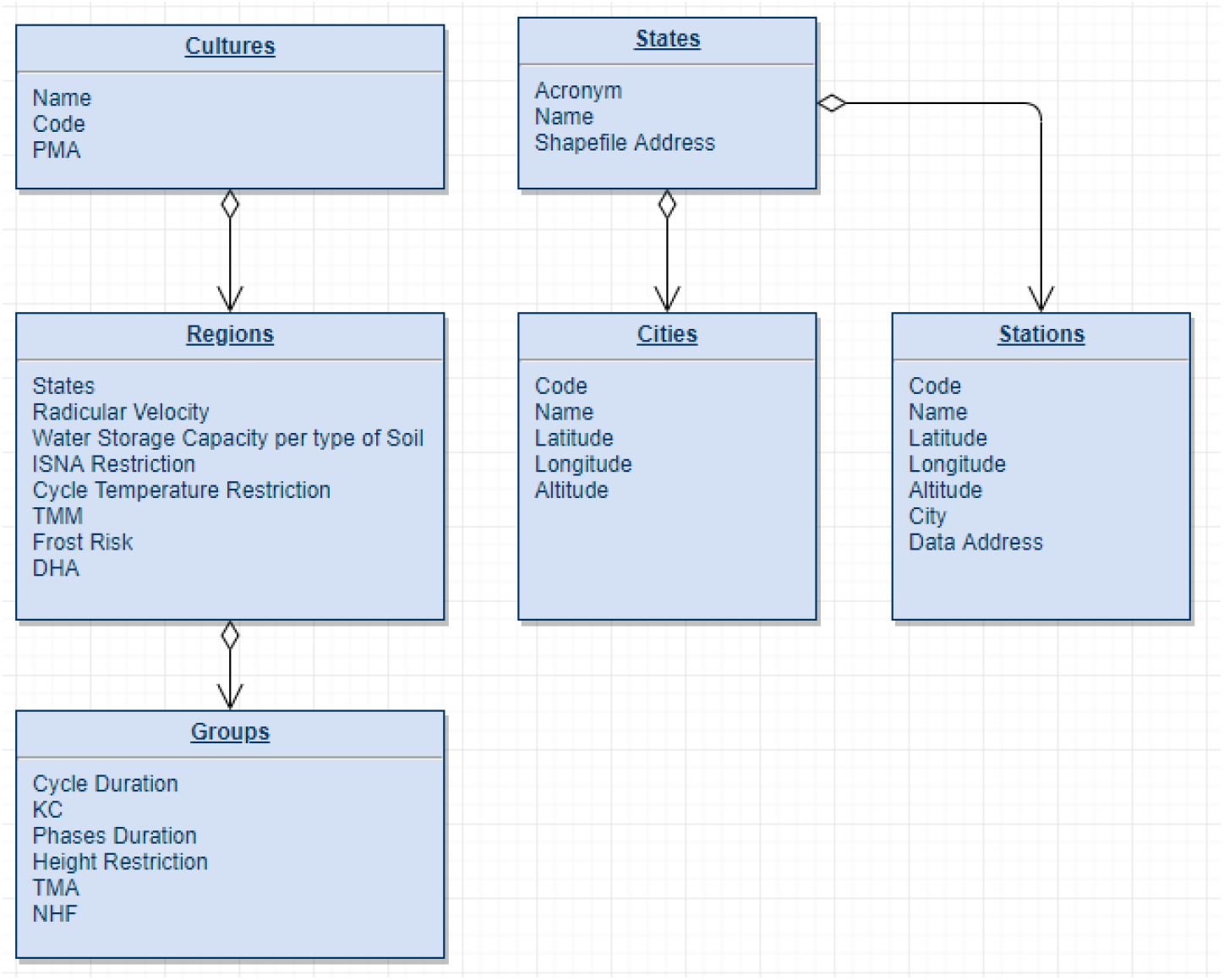
Schematic for the program behavior

To roll the analysis, the user needs to select the interest crop, type of soil (sand, silt or clay), plant group (advanced, normal or late), state and choose the climate database (Agritempo or EtaHadGEM2-ES current, optimist future scenario or pessimist future scenario). Then, the program pulls the necessary information that is associated to its group, soil, culture and state. Figure 1 shows how this data is linked on our culture/station database. Note that some requirements and information vary depending on culture or region or group.

Therefore, BRAMAZOS has the soil and group information, along with the culture requirements and a list of stations for the chosen state. As it can be seen on Figure 2, the stations are linked to their name, code, latitude, longitude, altitude, city, state and climatic data address.

The program chooses to receive data from either city or station, and this depends on how the selected database gathered climatic data. For each station or city, the water balance is calculated following the SARRA software (Baron et al. 1999). However, instead of SARRAMET pre-processing, we created our own tables and culture databases, along with stations climate data. We also calculated some variables using different methods – which will be explained below. In addition, a new method of zoning was developed for BRAMAZOS, using regional-related restrictions. This was important because SARRA was designed in France, which has different needs, and was adapted to analyze the climate changes over the development of cereals on tropical environments (Baron et al., 1999).

When creating a new simulation, SARRA has three main processes: carbon balance, water balance and phenology. Before that, it is necessary to define portion and soil characteristics, place, culture ecophysiological traces, cultivation practices, precipitation and temperature data for each station. In addition, the planting date should be chosen (Baron et al., 1999).

Comparing to that, BRAMAZOS already contains all the culture and stations data connected to the same database. The user only needs to choose the state, date and type of culture he wants to simulate. However, CEPAGRI’s developers have access to add new climate data, new stations and more cultures. In contrast, SARRA allows the user to add data by himself.

For BRAMAZOS software, this means that, instead of the user defining them, the surface runoff, root depth, mulch, radiation, relative humidity and storage are either calculated in the program or fixed values. On the plus side, BRAMAZOS’ users do not need deep agricultural knowledge, so it can be used by non-specialists, and not only agronomists. On the down side, those variables might have more accurate and local formulas based on the chosen state and country.

Discussing about calculation differences, in BRAMAZOS the monthly Potential Evapotranspiration (ETP) is calculated using Thornthwaite and Matter (1955) method, which is shown in the equations 2, 3, 4 and 5.

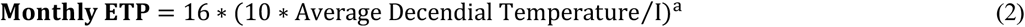

Therefore, the decennial ETP is:

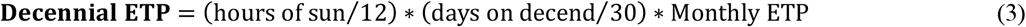

Also not related to SARRA, the program uses common zoning formulas, such as those below for Storage Variation (equation 4), Culture Parameter (Kc) and Culture Potential Evapotranspiration (EPC) (equation 5). The first one is mentioned by Camargo (1962):

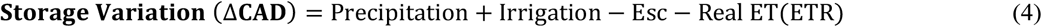

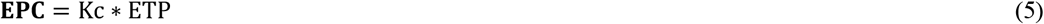

Where, Kc =proportion from decendial Kc table and days on current decend.

However, the Surface Flow (Esc) is calculated via Horton method (Horton, 1940), using the same parameters as SARRAZON (equation 6 and 7) and where the parameters is described in Table 1.

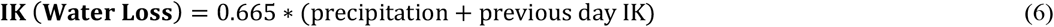

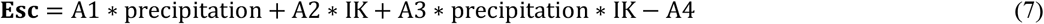

**Table 1 –.** Parameters to Surface Flow (Esc) calculation

| Soil Type | A1 | A2 | A3 | A4 |
| --- | --- | --- | --- | --- |
| 1 – Sand | 0.2 | 0.03 | 0.004 | 3 |
| 2 – Silt | 0.35 | 0.04 | 0.004 | 3 |
| 3 – Clay | 0.9 | 0.05 | 0.002 | 10 |

In addition, as stated before, BRAMAZOS’ water balance does follow main SARRA formulas and internal variables. They are listed in equations 8 to 14.

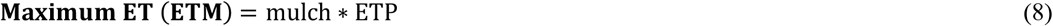

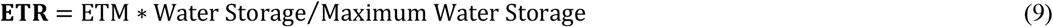

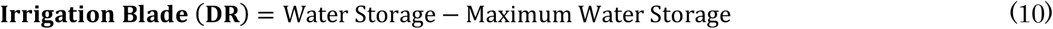

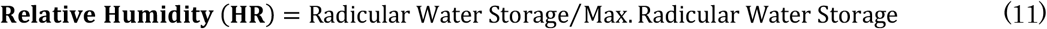

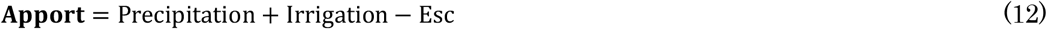

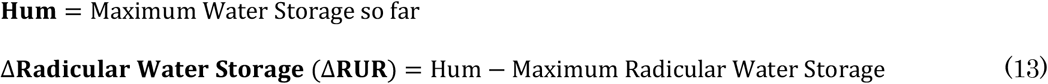

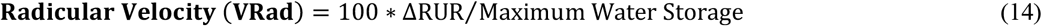

The value of Real Evapotranspiration (ETR) is calculated using Eagleman method (Eagleman, 1976), as suggested by SARRAZON and demonstrated in equation 15.

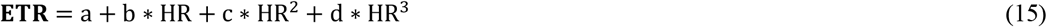

Where,

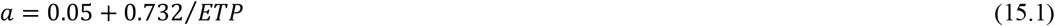

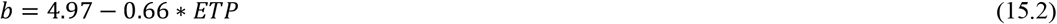

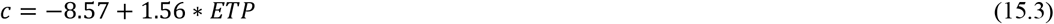

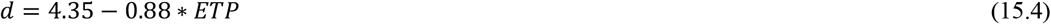

For calculating Normalized Water Balance (with monthly values), the equations were based on Thornthwait & Mather (1955) method simplified by Camargo (1962) for Ribeirão Preto area, in São Paulo/Brazil. The equations used are 16, 17, 18, 19, 20, 21 and 22.

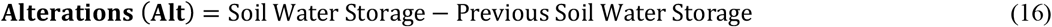

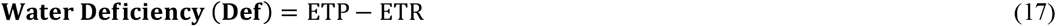

When Soil Water Storage (ARM) is equal to Maximum Water Storage (CAD):

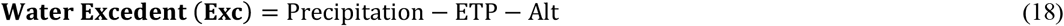

For ETP higher than precipitation:

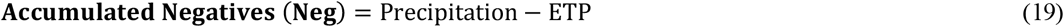

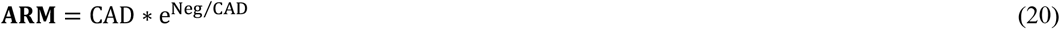

For precipitation higher than ETP:

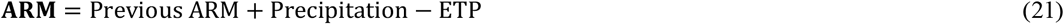

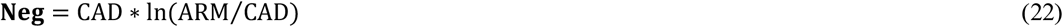

#### b) Implementing Restrictions

After calculating the water balance variables, two tables are returned: one with daily values of ETP, Esc, Apport, Kc, ETR, Hum, DR, HR, Radicular Velocity, Maximum RUR, RUR, EPC, ETR, ETM, Minimum Temperature, Mean Temperature, Maximum Temperature, Precipitation, etc. The other has the normalized monthly water balance values: Precipitation, ETP, ARM, Neg, Alt, ETR, Def and Exc.

The final result is calculated from these tables and the restriction list for the selected culture. For each restriction, it is given a positive result if at least 80% of the year’s data is within selected range.

A new table, which is the only one who will be exported from the Simulation script, is created. The first column represents a percentage of restrictions that had positive result, and the columns after that give the percentage of years that were within the low-risk range for each restriction.

For instance, Persimmon, Fig and Papaya were chosen to be analyzed. Their requirements are listed on Table 2, 3 and 4. The PMA variable is calculated through the Normalized Water Balance (BHN) table. This is done by creating a list with the sum of the 12 monthly Precipitation values for each year in the table. A percentage of the years where the sum is above 1000 (for Persimmon) or 1200 (for Fig) is the result for PMA. Therefore, if this percentage is above 80%, then to the final result it will be added ¼ (for Persimmon) or 1/3 (for Fig), equivalent to 1 positive restriction.

**Table 2.** Persimmon requirements to run Bramazos software

| Variables |  | Unit | Optimal | Suboptimal | Unsuitable |
| --- | --- | --- | --- | --- | --- |
| **PMA** | All groups | mm | >1000 | - | <1000 |
| **TMA** | Group I | °C | >17 and <22 | >14 and <22 | <14 or >22 |
|  | Group II | °C | >15 and <18 | >14 and <28 | <14 or >28 |
| **Frost risk** | All groups | % | <20 | - | >20 |
| **NHF** | Group I | h | >100 | - | <100 |
|  | Group II | h | >300 | - | <300 |

**Table 3.** Fig requirements to run Bramazos software

| Variables | Unit | Optimal | Suboptimal | Unsuitable |
| --- | --- | --- | --- | --- |
| **PMA** | mm | >1200 | - | <1200 |
| **TMA** | °C | >20 and <25 | >15 and <36 | <15 or >36 |
| **Altitude** | m | >700 | - | <700 |

**Table 4.**
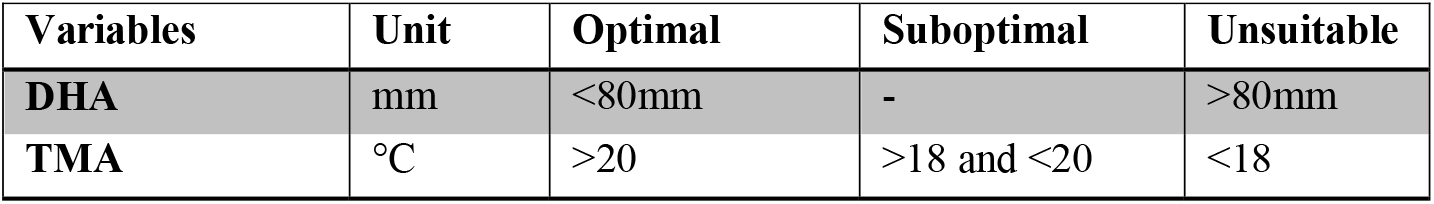
Papaya requirements requirements to run Bramazos software

The TMA value is calculated similarly to the PMA. The main difference is that TMA has one extra class (Suboptimal result). To the code, this means that if the temperature is within suboptimal parameters, the result is 50% positive, and if it is also within optimal parameters, more 50% is added to the result. Of course, the 80% of the year’s restriction has to be satisfied. In addition, the same logic is applied to the final result: suboptimal result for TMA adds 1/8 (for Persimmon) or 1/6 (for Fig) and optimal result adds more 1/8 or 1/6.

Frost Risk is calculated using the formula 23.

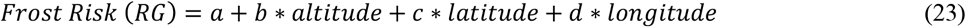

Where altitude is in meters and latitude and longitude are in minutes. However, a, b, c and d depend on the temperature reference, as specified on Table 5. If the result is below the requirement, then ¼ is added to the Persimmon result.

**Table 5.** Temperature reference to calculate Frost Risk (RG)

| Temperature | a | b | c | d |
| --- | --- | --- | --- | --- |
| 0°C | -246.64 | 0.0568 | 0.1217 | 0.0221 |
| 1°C | -325.13 | 0.0606 | 0.1572 | 0.0348 |
| 2°C | -374.45 | 0.0616 | 0.1808 | 0.0446 |

In order to implement altitude restriction, the result 100% is given if the station or city altitude is within 100m of the optimal restriction and 0% result if otherwise. This counts as 1/3 of the Fig final result.

To analyze NHF, the formula (24) was used (for temperatures below 13°C) for each year within climatic data.

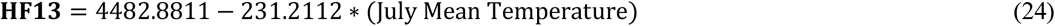

Therefore, the percentage of years can be calculated and ¼ is added to the Persimmon final result if positives are above 80%. The results were evaluated according to Junior et al. (1979), where the formula above is referenced.

When the user chooses to analyze data and fills the fields, a new page will be open with several tabs. The first one contains a map with the percentage of rightful criterias (final result) and for each restriction there is a new tab with a corresponding map. Because of that, it is possible for the user to see the suitability of the chosen area for each variable, and therefore understand the reasons why the result was positive or not.

Inside each tab, there is a map with four coordinate lines, a wind rose, scale, subtitles and five color zones according to the result (0-20%, 20-40%, 40-60%, 60-80%, 80-100%). In addition, there is a title with the culture name and planting date and on the lower side of the page there is information about the chosen database, group and soil.

### 2.3 Case of study: Fruits and simulations

A case of study is presented to illustrate the usage of the software to climate risk analyses to two species of temperature fruits - Persimmonn (*Diospyros kaki* L.) and Fig (*Ficus carica* L.) - and one tropical fruit - Brazilian Papaya (*Carica Papaya* L.) - in their cultivation area in São Paulo State, southeast of Brazil.

#### a) Study area

The software is being adapted to run in all over Brazilian soil and climate conditions. The experiments simulation presented in this study were carried out at São Paulo state, southeast of Brazil (Figure 3). According to Herrera et al. (2017), 11.7% of the Brazilian family farmers are located in this region, being the second leader in gross production value (GPV) in the country. This state was choose because the family farmers do not present high dependence on irrigation to increase productivity. Additionally, São Paulo state accommodate the “Fruit Circuit” region, which is an important stallholder’s area, responsible for fruits production, especially grape, strawberry, peach, guava, plum, persimmon, acerola and fig.

**Figure 3 –.**
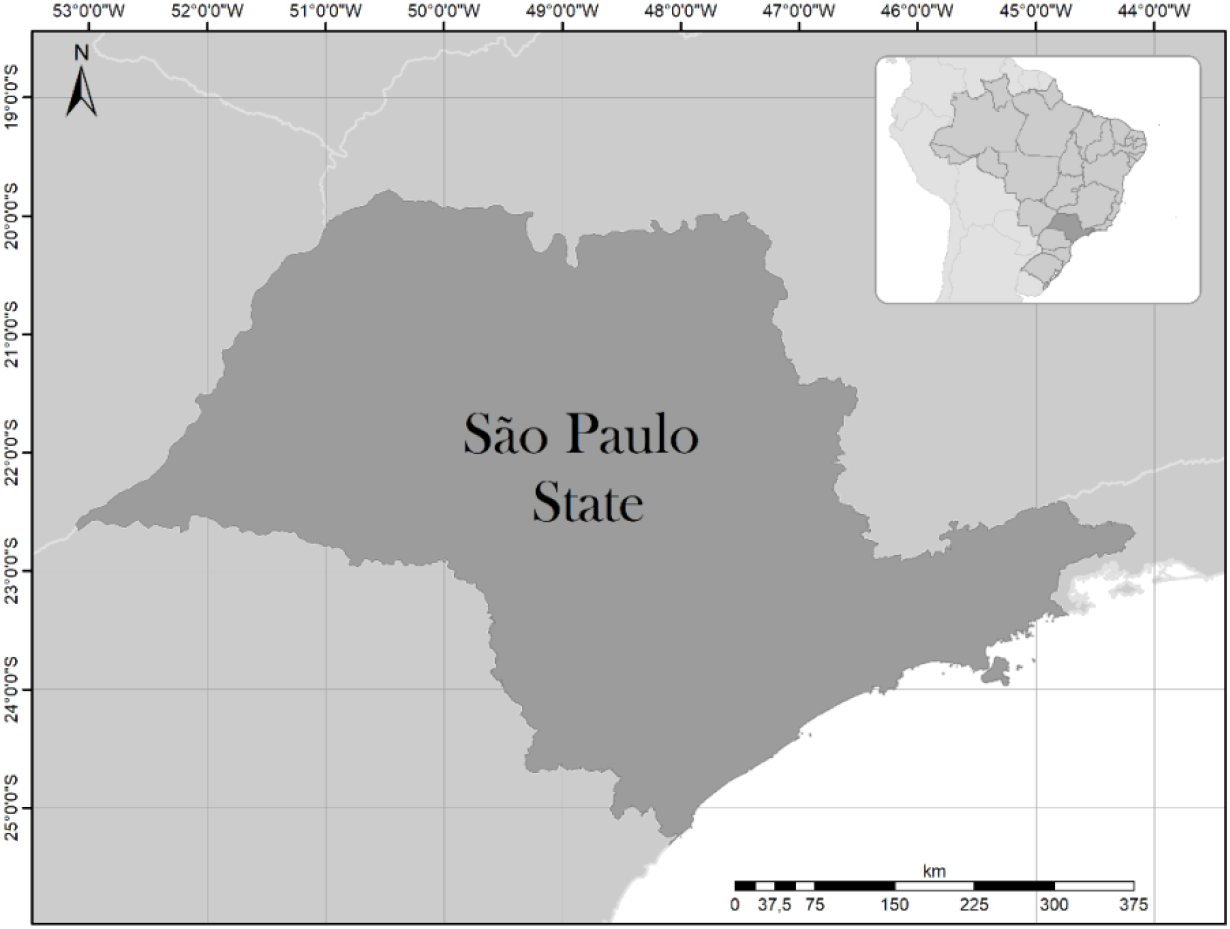
Study area: São Paulo State, southeast of Brazil

#### b) Fruits and Simulation

Brazil is the third most important fruit producer in the world (Retamales et al., 2011) and, according to the Brazilian Yearbook of Fruticulture (2018) the top fruit crop in the country yielded around 43.5 million tons of fresh fruit in 2017, in which 65% is consumed inside the country and 35% are destined to exportations. Fruits produced in Brazil supply consumers of about 100 countries in the world generating opportunities for small-scale business and smallholders. According to the Brazilian Fruit Growers and Exporters Association, in 2017 the total fruit exports amounted to 878.4 thousand tons and brought in US$ 946.8 million, demonstrating that Brazilian fruits are very popular with the international consumers. The fruit production is increasing in all Brazilian states but São Paulo has been considered, in the last years, as the biggest national producer. The fruit farmers in São Paulo harvested 15.9 million tons in 2015, followed by Bahia (4.9 million tons), Minas Gerais (3.1 million tons) and Rio Grande do Sul (2.7 million tons) (Brazilian Yearbook of Fruticulture, 2017).

The Brazilian fruit chain is quite dependent on the climatic conditions. In 2016, for example, the Brazilian Yearbook of Fruticulture published that the national fruit farming business reaped smaller than expected crops over the past years due to unfavorable climate conditions, such as prolonged drought in the Northeast; and rainfall, hailstorms and other climate related problems in the South-east and South of the Country. According to the Yearbook registers, the adverse weather conditions had also a say in the decline of Brazilian fresh fruit exports in 2016. The Brazilian Institute of Geography and Statistics (IBGE) related that the most cultivated fruit trees grown in Brazil reached an output of 40.953 million tons of fresh fruit in 2015, decreasing of 1.7 million tons of fresh fruit, in comparison to the 42.6million tons registered in 2014.

To run the simulation, we choose two examples of temperate fruits (Persimmon and Fig) and one temperate fruit (papaya).

Persimmon (Diospyros kaki L.) is a deciduous fruit tree originally from China and was introduced in São Paulo, Brazil, in the end of XIX century (Martins and Pereira 1989). The main world producer is China, followed by Japan, South Korea and Brazil (FAO, 2007). According to Martins and Pereira (1989), the persimmon is typically subtropical fruit and, despite the deciduous habit, is able to adapt to different climate and soil conditions. This adaptability allowed its distribution in different states of Brazil, mainly in the south and southeast regions, where it became a very popular fruit. Although persimmon is not a fruit with enormous prominence, the fruit crop has grown the most in planting and production area in the last 20 years in Brazil, and São Paulo State is highlighted in this production. Specified persimmon climate requirements are described in table 5.

Following the Persimmon tendency, the fig (*Ficus carica* L.) fruit production is increasing in Brazil and, since 1995, according to IBGE, the harvesting area increase in average, 38% per year. In São Paulo state, the growth rate is, in average, 30% per year. Originally, from Asia and Syria, Fig is one of the oldest cultivated planta in the world (Veberic and Mikulic-Petkovsek, 2016) and can be consumed fresh, dried, canned or candied (Sinha, 2003). The fruit also has medicinal properties (Ercisli et al., 2012), which improve demand. Fig does not require a very rich soil for cultivation and normally is considered as drought tolerant crop (Sinha, 2003). As the Persimmon case, Fig is also be able to adapt to different climate conditions and is normally cultivated in warm and dry climate (Veberic et al., 2008). Specified Fig climate requirements are described in table 6.

**Table 6.**
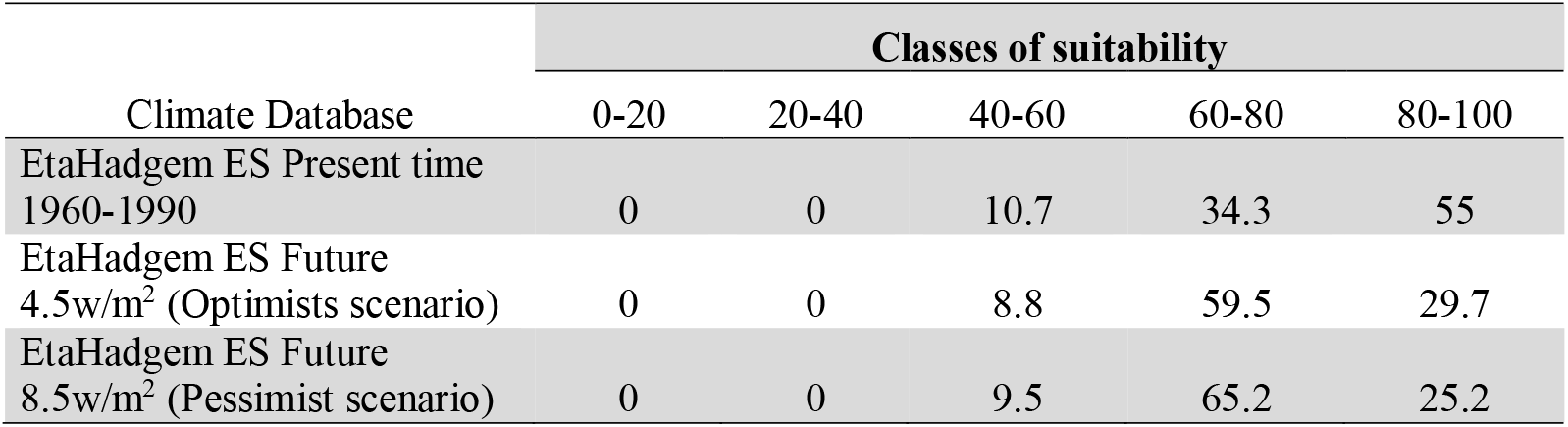
Surface of São Paulo are (in percentage) classified by level of suitability for Persimmon production in present time and future climate change scenarios, using the climate regional model EtaHadgem ES

Brazilian Papaya (*Carica Papaya* L) is an important horticultural crop to tropical and subtropical regions and Brazil is the second most important producer in the world, following India (Dantas et al., 2018). In terms of production value, Papaya occupies the fourth place following orange, bananas and grapes (Carvalho et al., 2014). The exact center of origin is still unknown, but it is considered that Papaia is probably native from Mexico and Central America, being very sensitive to frost and chilling (Morton, 1987). The cultivation needs generous rainfall (or irrigation), but the soil must present good drainage.

Given these Persimmon, Fig and Papaya characteristics and, knowing the importance of these crops for family farming, it is important to support smallholders to manage their climate risk on the production area using a simple and robust tool. In this context, three simulations were performed in this study:

a. First, we did a simulation with EtaHadgem ES to the current time (using 1960-1990 time series data)
b. Second, we design a simulation with EtaHadgem ES future climate change, in an optimist scenario (which consider low emissions of greenhouse’s gases – 4.5M/m^2^)
c. Third, we performed a simulation with EtaHadgem ES future climate change pessimist scenario (which consider high emissions of greenhouse’s gases – 8.5M/m^2^)

## 3. Results and Discussion

### 3.1 General BRAMAZO Overview

Figure 4 presents the software interface, which is responsible for receiving user information allow the command to corresponding simulations. The required parameters are loaded by the database. When the user starts the program, some specific and basic information is required such as:

- crop: the user can choose between the crops that are implemented: annual crops (cotton, rice, beans, corn and soybean) or perennial crops (banana, coffee, pineapple, persimmon, citrus, fig, papaya, passion fruit, nectarine, peach, mango and manioc.
- group, which is the planting group, related to the cycle of the crop. There are three different group that the user can choose: precocious, normal and late.
- soil type, choosing if the soil is sand, silty or clay;
- State: that is the federation unit, where the farmer can choose the local where he wants to plant the crop;
- climate database, where it is possible to choose between meteorological stations database or climate model ETAHadgem ES in three possible scenarios: actual (1960-1990) or future scenarios, being one of them the optimist and the other is the pessimist;
- planting date: which is the date when it is planning (sowing) to start the cultivation;
- interpolation: if the user wants to see the results with or without interpolation
- option: if the user wants the result in map or tables

**Figure 4.**
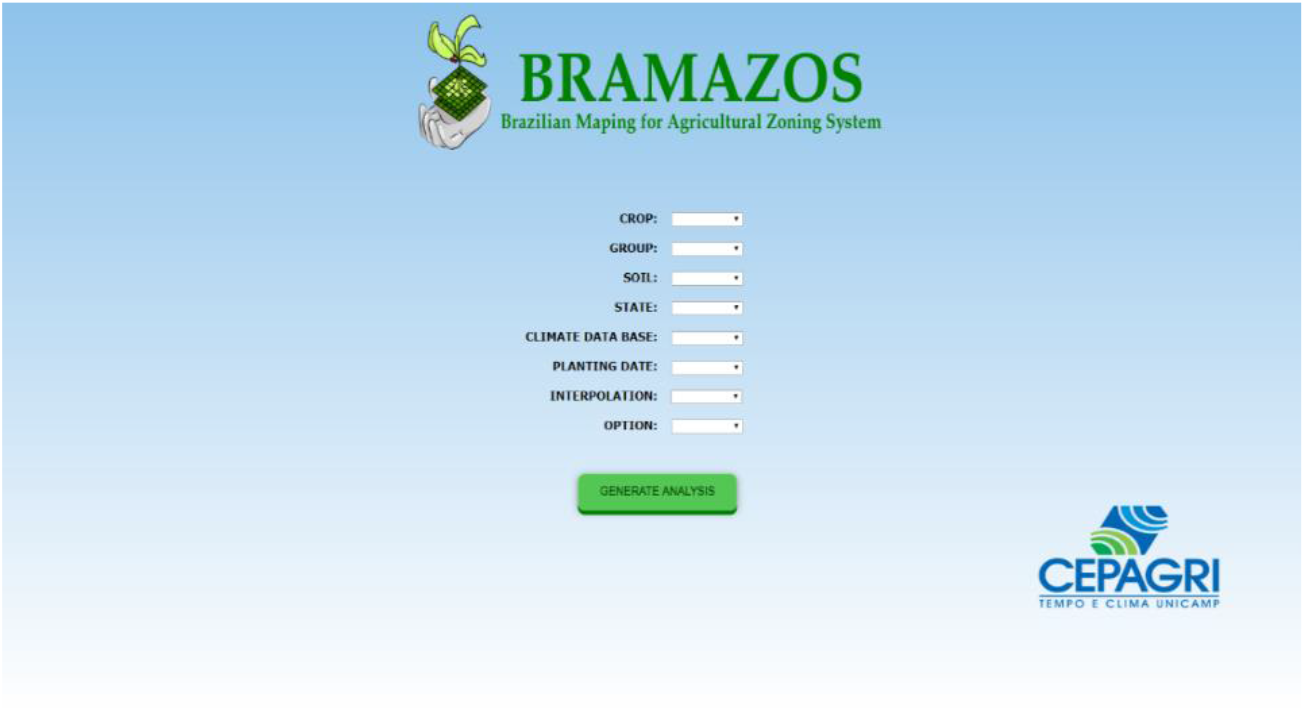
BRAMAZOS Interface

After that, the user can run the simulation, choosing the “generate analyses bottom”. The results, in map format are presented in figure 5. The user is able to choose the variable to visualize the result; therefore, the farmer can understand what is the main restriction, eg. High temperature risk, frost possibilities, precipitation limitation (water deficit), or geographical limitation such as altitude. Using this information the farmer is also capable to manage this limitation with agriculture field management.

**Figure 5 –.**
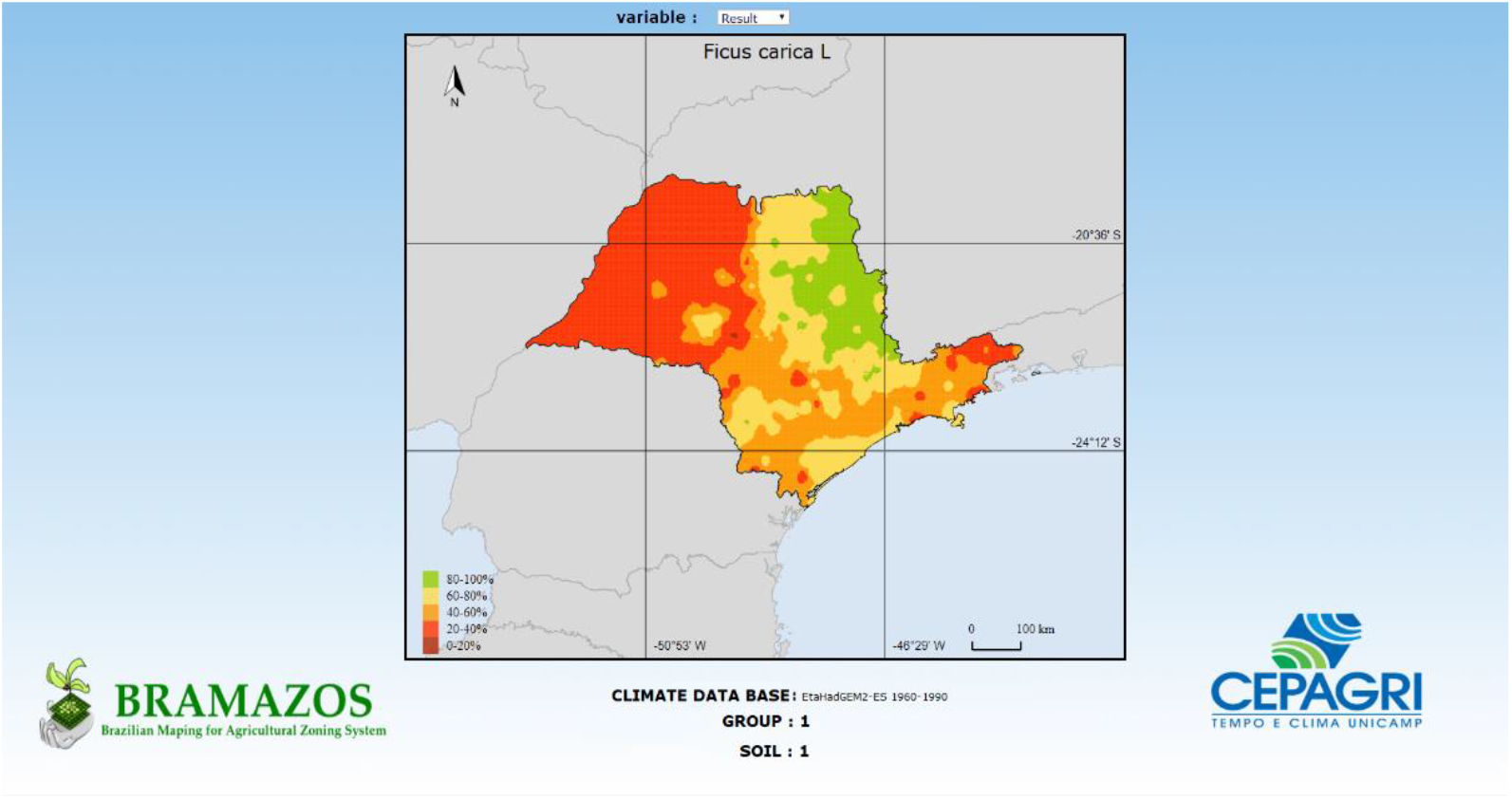
Result of the simulation using the “map option” answer

### 3.2 Persimmon Climate Risk Analyses

Figure 6, 7 and 8 presents the results from BRAMAZOS software, using EtaHadgem ES Present time Database (1960-1990) (Fig 6), EtaHadgem ES Future Optimist scenario (Fig 7) and EtaHadgem ES Future Pessimist Scenario (Fig 8). Table 6 summarizes the results presented in Figures. In current situation, 55% of São Paulo landscape, presented precipitation, temperature, chilling requirement and frost risk suitable for persimmon cultivation, in, at least, 80% of the evaluated climatic series. 34.3% of the total area was classified in the second class of suitability (with three meteorological variables satisfactory) and 10.7% presented two meteorological variables satisfactory for this cultivation. The main meteorological variable that behaved as a limiting factor to cultivation was temperature, especially in the chilling requirements and annual average temperature risk. 45% of the total area was limited by high temperatures.

**Figure 6 –.**
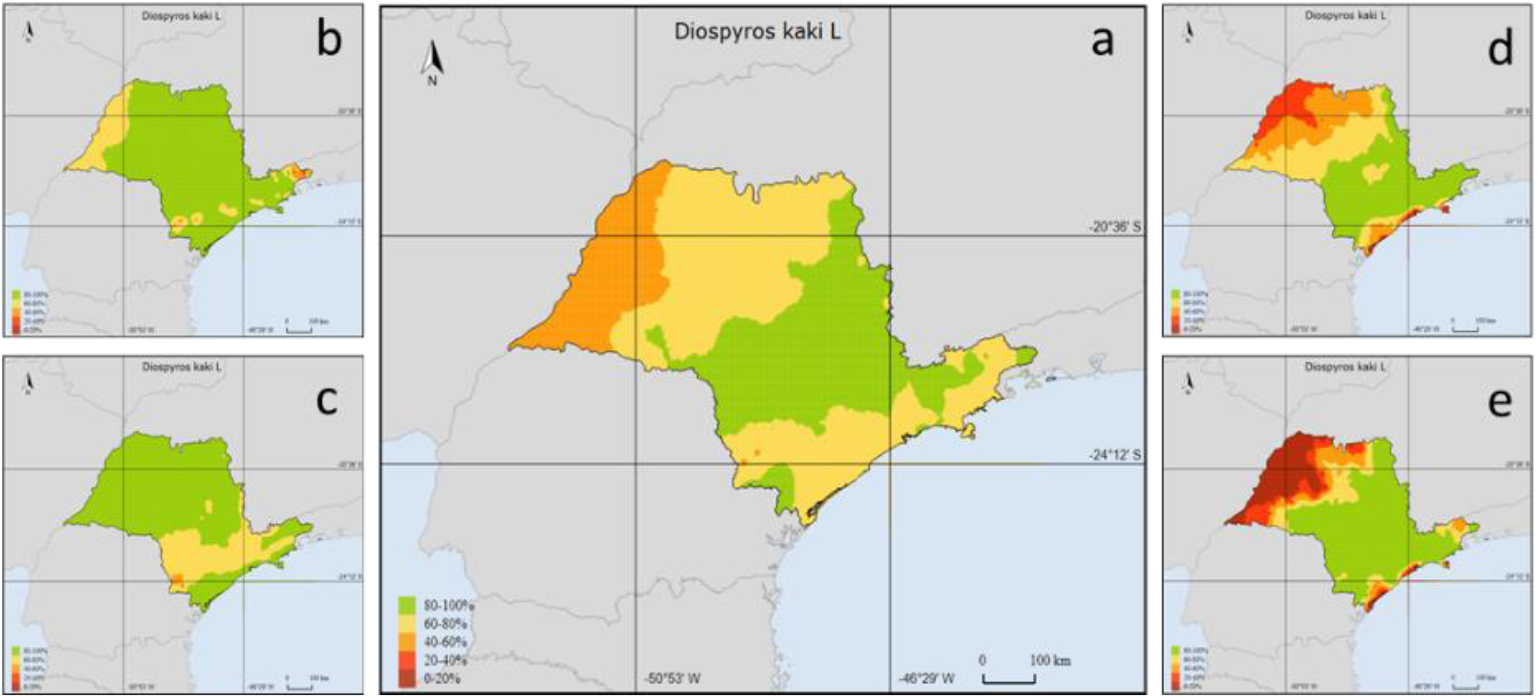
Suitable area in São Paulo State, Brazil, for Persimmon cultivating using EtaHadgem ES Present time Data base (1960-1990). In a: result with all variables, b: precipitation restriction; c: frost risk; d: chilling requirements and e: annual average temperature risk

**Figure 7 –.**
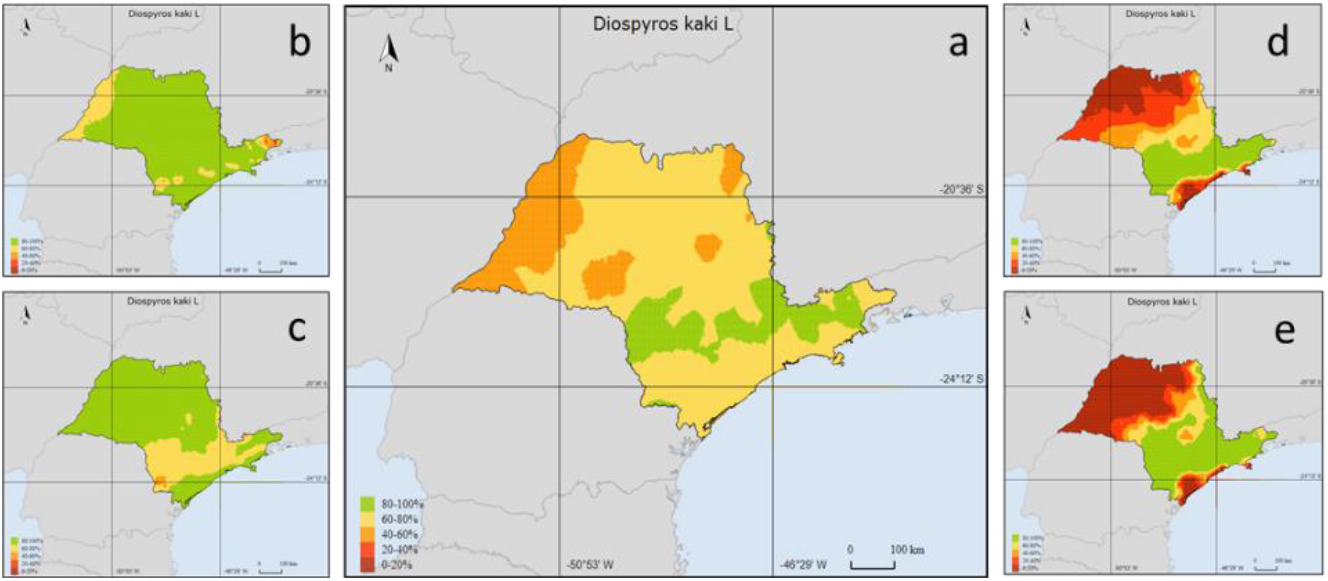
Suitable area in São Paulo State, Brazil, for Persimmon cultivating using EtaHadgem ES Future 4.5w/m2 Optimist Data base. In a: result with all variables, b: precipitation restriction; c: frost risk; d: chilling requirements and e: annual average temperature risk

**Figure 8 –.**
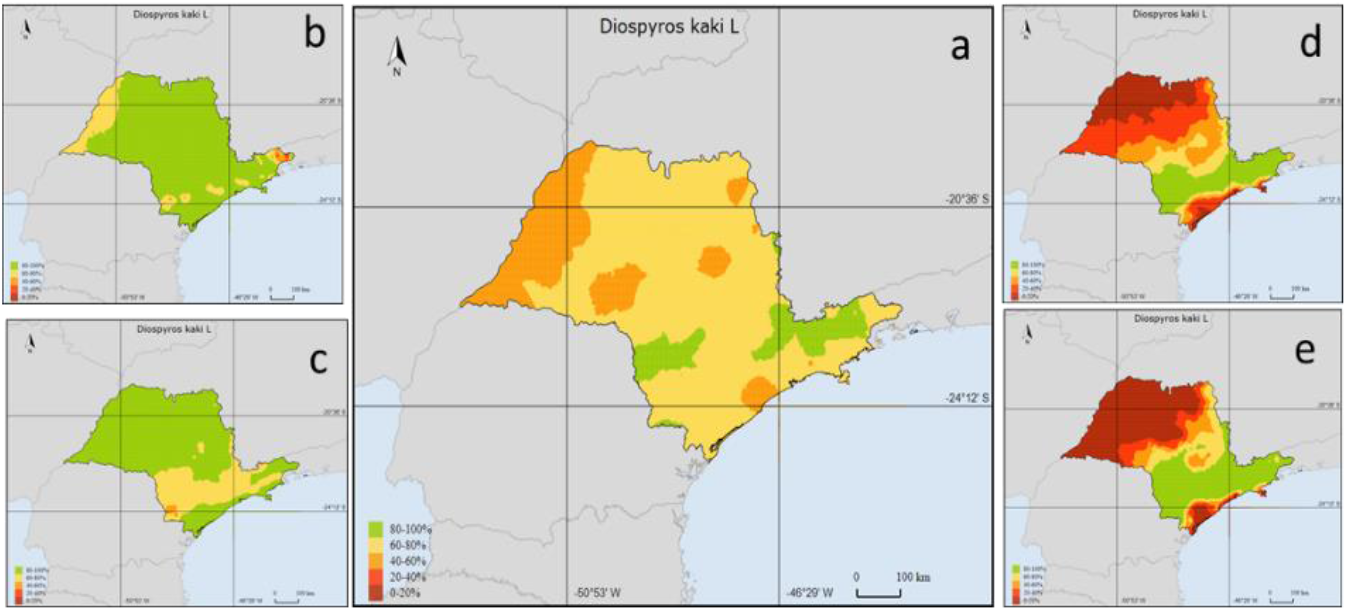
Suitable area in São Paulo State, Brazil, for Persimmon cultivating using EtaHadgem ES Future 8.5w/m2 Pessimist Data base. In (a): result with all variables, (b): precipitation restriction; (c): frost risk; (d): chilling requirements and (e): annual average temperature risk

An important point to consider in Bramazos system is that the software is able to indicate which meteorological variable is behaving as a limiting factor, allowing smallholders to manage the climate risk. Understand and identify the suitability class and the climate related risk are important for improve adaptation option activities based on scientific knowledge. Taking precipitation as an example, if this meteorological variable is the limiting factor, the farmer may choose use irrigation system to compensate the rainfall limitation.

Significant changes in the suitability zone distribution were visible in the projected scenarios of climate change, were the suitable area (class of suitability 80-100%) decreases 25.3% and 29.8% for the optimist and pessimist scenarios respectively. The increase in temperature, which is a condition not current seen in the west of São Paulo, appears to be the major limiting factor for the future cultivation. The exposure to chilling conditions deficiency due to the increase in the annual average temperature tend to decrease the suitable area for persimmon cultivation, resulting in a displacement of the suitable zone, which will be concentrated mainly in higher elevation regions and in the south of the State, where the temperature is naturally lower. Similar results were found in Brazil to other cultivation, as coffee Arabica (Zullo Junior et al., 2011), which could change the production area to the extreme south of the country and to higher elevation regions, with colder and moderate climate.

Although Persimmon can be cultivated in a wide range of subtropical and warm temperate climate, high temperatures could affect the dormancy period (Mowat et al., 1995), accelerate the flowering process (George et al., 1997) the development of the flower embryo (Fukui et al., 1990) and the fruit set rates (Zilkah et al., 2013), prejudicing the production and quality of the fruit. Case studies in peach trees (Kozai et al., 2004) and to grapevine in Bordeaux (Jones and Davis 2000) found similar results, suggesting that warming day make the crop cycle precocious and the season duration shorter. In the grapevines, the impact of the decreasing season is the harvest logistic and the wine quality (Webb et al., 2007) and these impacts could be similar to the persimmon, affecting the logistic and commercialization as the fruit quality.

Zilkah et al. (2013) studying the effect of high temperature on Persimmon showed decline of fruit set when the trees were exposed to high temperatures during the flower development period and Lazar (2008) found a positive correlation between fruit deformation rate and fruit drop on the tree. These studies together suggest that high temperature is probably related to fruit decline and deformation. High temperatures, associated to water stress, during the floral differentiation can also be unfavourable, inducing abnormal flowers (George et al. 1997)

To mitigate the effects of high temperature in the Persimmon cultivation, Zilkah et al (2013) suggest protective treatments, such as shading. Zilkah et al (2013) showed a significant reduction in fruitlet, branches and leaf temperature in the shading cultivation. Similar results were also found in cultivation that is sensitive to high temperatures, such as the coffee arabica (Lin 2007, Pinto et al. 2008). These studies, recommended shading as an important technique to adapt the cultivation to climate changing scenarios. In addition to reduce high temperatures, shade management technique, is an important option to mitigate the greenhouses gases, because this kind of cultivation can stock more atmospheric carbon than the cultivation alone (Coltri et al. 2015). Other adaptation management suggested by Zilkah et al (2013) to Persimmon, is evaporative cooling, sprinkling water and cooling the fertility organs avoiding fruitlet dropping. The study showed a dropping reduction approximately 45% using this management technique.

### 3.3 Fig Climate Risk Analyses

Figure 9, 10 and 11 presents the results from BRAMAZOS software, using EtaHadgem ES Present time Database (1960-1990) (Figure 9), EtaHadgem ES Future Optimist scenario (Figure 10) and EtaHadgem ES Future Pessimist Scenario (Figure 11). Table 7 summarizes the results presented in Figures. In current situation, 20.9% of São Paulo landscape presented class of suitability 80-100%, indicating that precipitation, annual average temperature, and altitude are satisfactory for the cultivation. São Paulo state presents a significant restriction due to the altitude, as we can verify in figure 9 (d). The suitable area with all the meteorological variables satisfactory to the fig cultivation may present a small decrease in future scenarios of climate change, reducing less than 1% of the suitable area.

**Figure 9 –.**
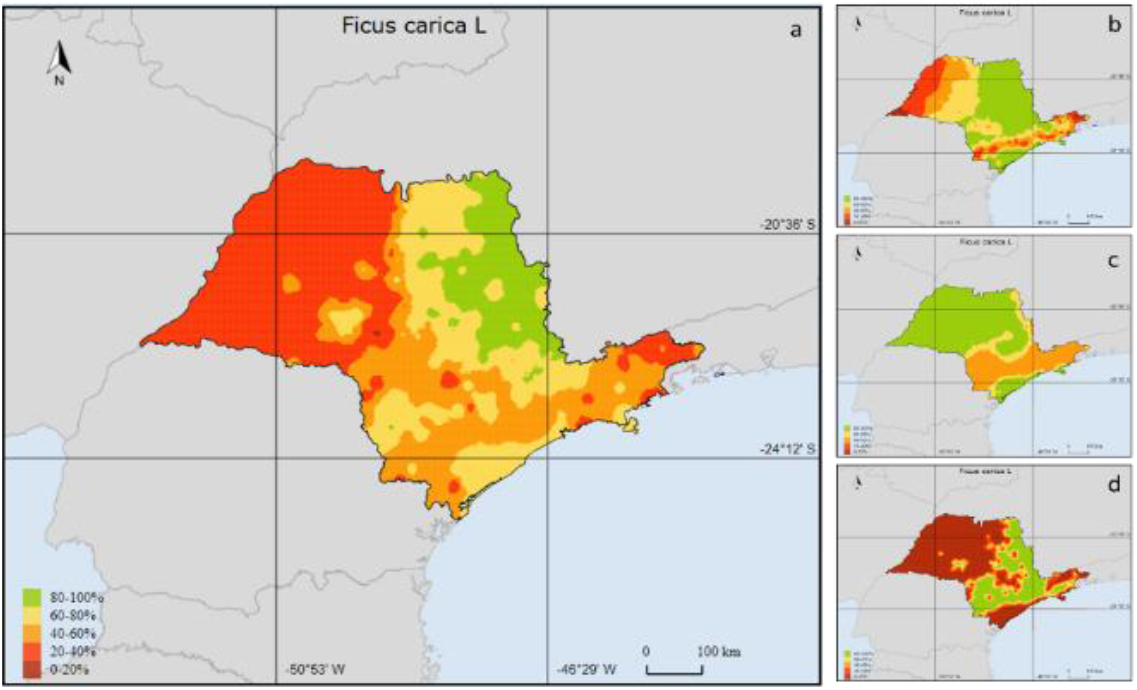
Suitable area in São Paulo State, Brazil, for Fig cultivating using EtaHadgem ES 1960-1990 Data base. In (a): result with all variables, (b) precipitation restriction (c) annual average temperature risk and (d) Altitude restriction for Fig cultivation

**Figure 10 –.**
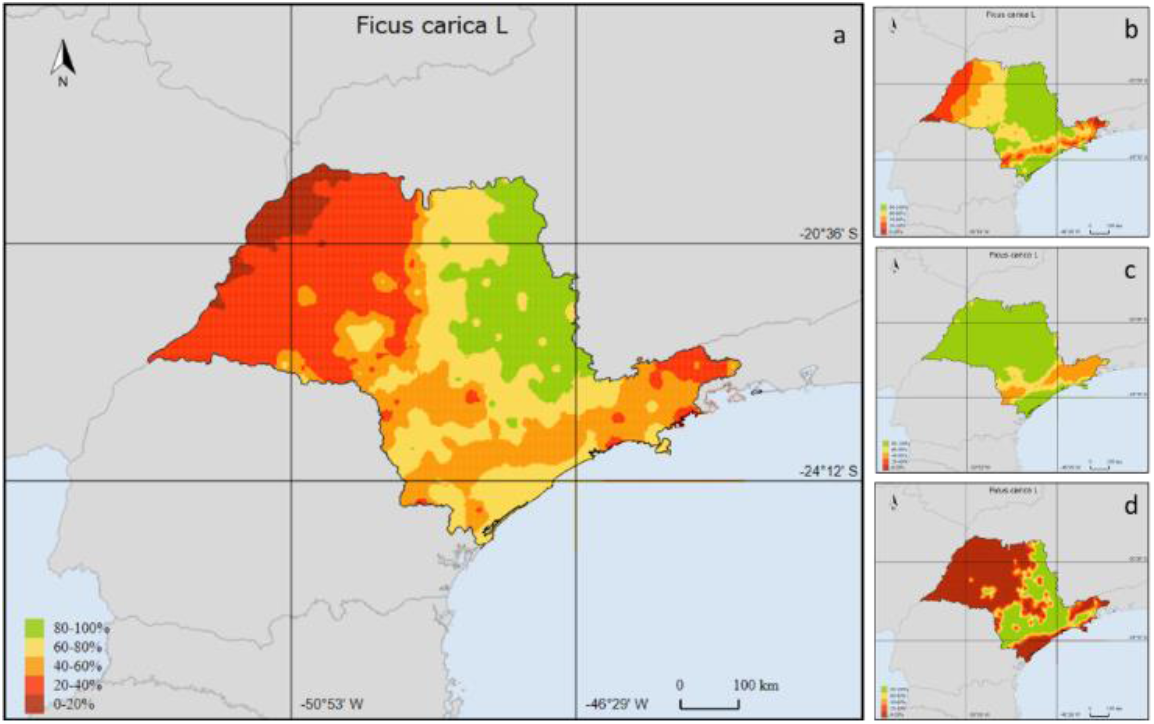
Suitable area in São Paulo State, Brazil, for Fig cultivating using EtaHadgem ES future optimist scenario. In (a): result with all variables, (b) precipitation restriction and (c) annual average temperature risk; (d) Altitude restriction for Fig cultivation

**Figure 11 –.**
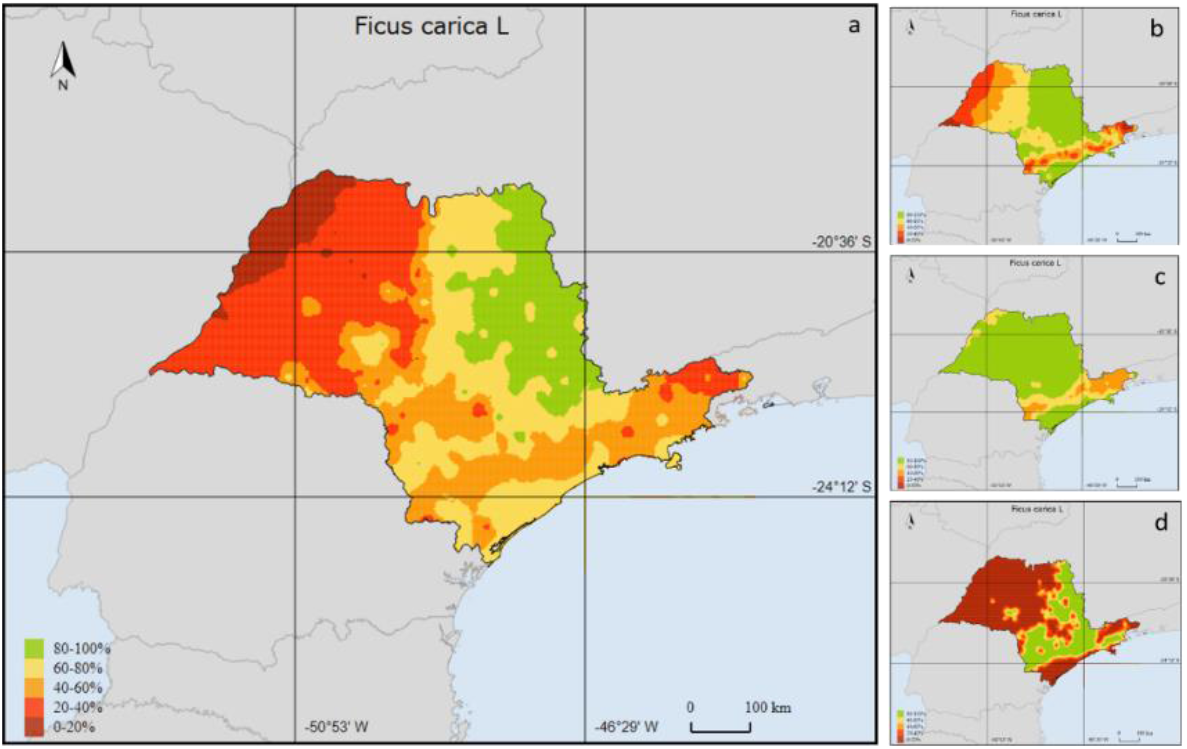
Suitable area in São Paulo State, Brazil, for Fig cultivating using EtaHadgem ES future pessimist scenario. In (a): result with all variables, (b) precipitation restriction and (c) annual average temperature risk, (d) Altitude restriction for Fig cultivation

**Table 7.** Surface of São Paulo are (in percentage) classified by level of suitability for Fig production in present time and future climate change scenarios, using the climate regional model EtaHadgem ES

| Climate Database | Classes of suitability |  |  |  |  |
| --- | --- | --- | --- | --- | --- |
|  | 0-20 | 20-40 | 40-60 | 60-80 | 80-100 |
| EtaHadgem ES Present time 1960-1990 | 4.7 | 35.9 | 19.7 | 18.6 | 20.9 |
| EtaHadgem ES Future 4.5w/m<sup>2</sup> (Optimists scenario) | 9.5 | 30.0 | 14.5 | 24.0 | 20.0 |
| EtaHadgem ES Future 8.5w/m<sup>2</sup> (Pessimist scenario) | 11.1 | 28.0 | 15.2 | 25.0 | 20.6 |

In future scenarios of climate change, although the suitable area for planting does not present a significant change, we can verify an increase of area with greater climatic restrictions. This, however, will not affect the fig culture because these areas were already restricted by altitude. In the suitable area, as in the persimmon case, annual average temperature appears to be the most affected variable in future climate change scenarios and what may be responsible by reducing fig cultivation area. Therefore, the results indicate that Fig, as it is planted and cultivated nowadays, could be a key crop to fruits smallholders in the futures scenarios of climate change, where the suitable zone tends to be not so affected.

### 3.4 Papaya Climate Risk Analyses

Figure 12, 13 and 14 presents the results from BRAMAZOS software, using EtaHadgem ES Present time Database (1960-1990) (Figure 12), EtaHadgem ES Future Optimist scenario (Figure 13) and EtaHadgem ES Future Pessimist Scenario (Figure 14). Table 8 summarizes the results presented in Figures. In current situation, Papaya has 32.7% of São Paulo landscape are suitable for its cultivation, with temperature and annual water deficiency satisfactory. In the projected future climate change scenarios, Papaya could lose around 9% of the suitable area, mainly due to increase of annual water deficiency. There are no significant difference between the optimist and the pessimist scenarios. Contrary to the fig and persimmon simulations, that presented to be more sensitive to the thermal requirements, Papaya appears to be more sensitive to the precipitation decrease in future projected climate change scenarios.

**Figure 12 –.**
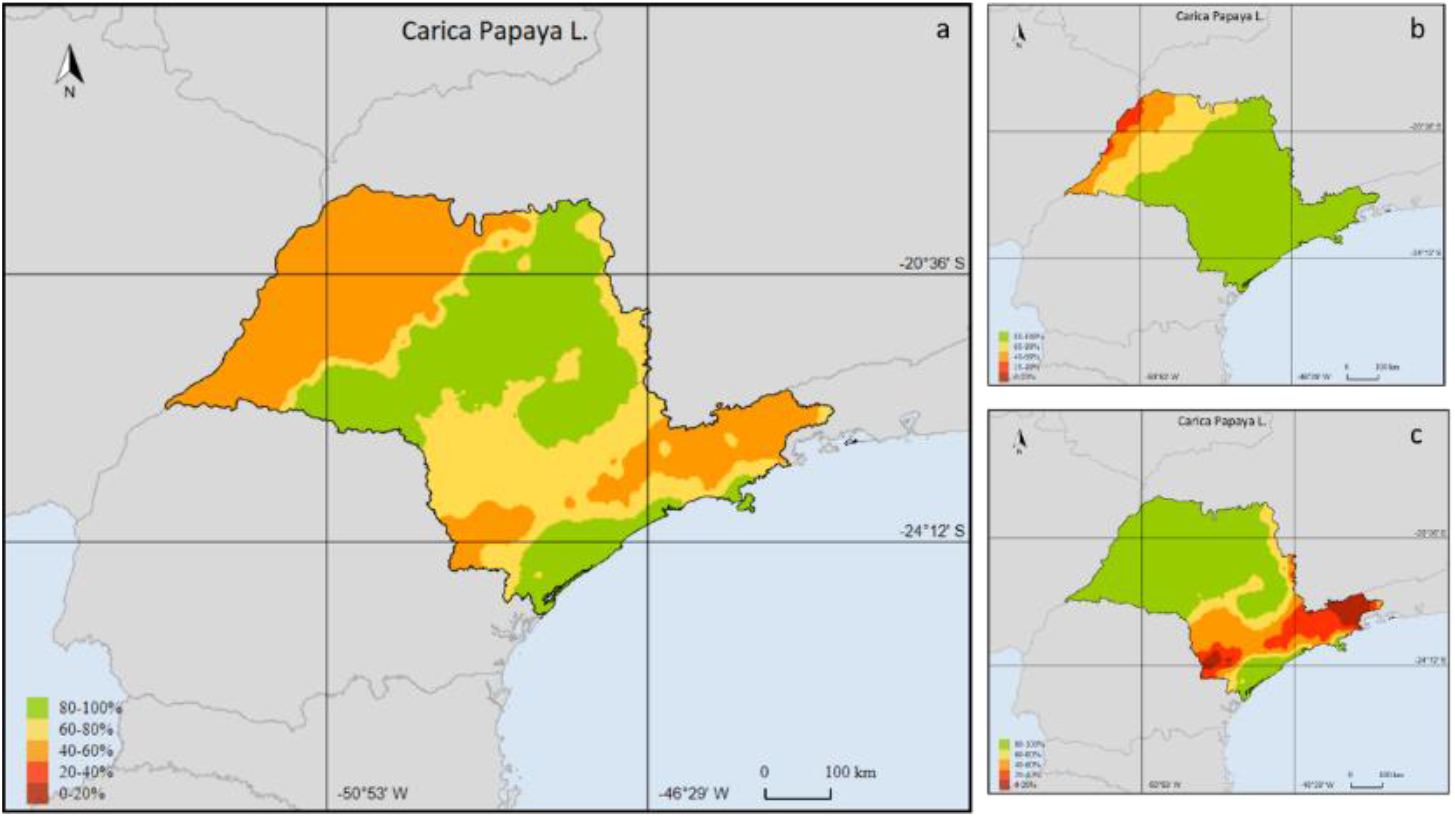
Suitable area in São Paulo State, Brazil, for Papaya cultivating using EtaHadgem ES 1960-1990 Data base. In (a): result with all variables, (b) annual water deficiency and (c) annual average temperature risk

**Figure 13 –.**
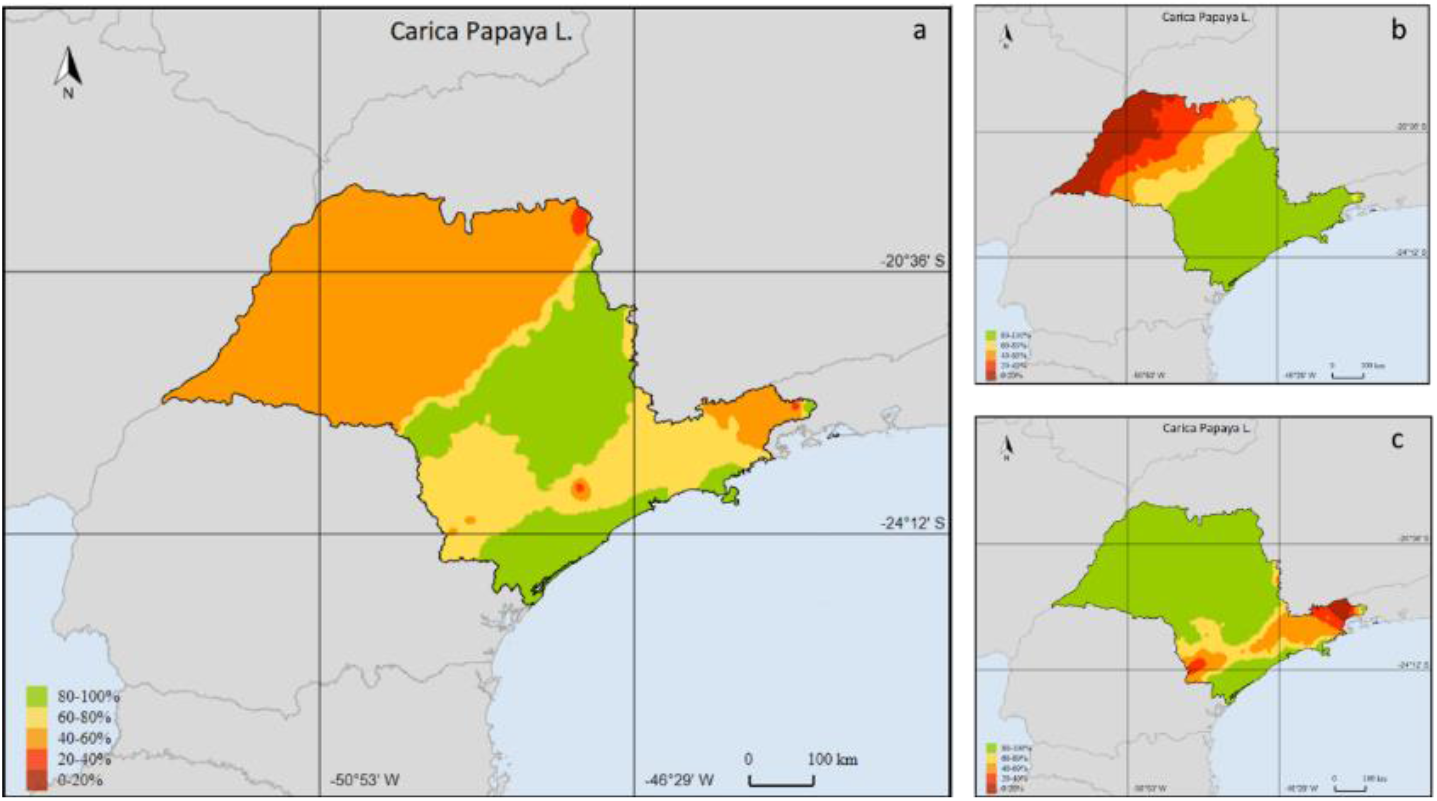
Suitable area in São Paulo State, Brazil, for Papaya cultivating using EtaHadgem ES future optimist scenario. In (a): result with all variables, (b) annual water deficiency and (c) annual average temperature risk

**Figure 14 –.**
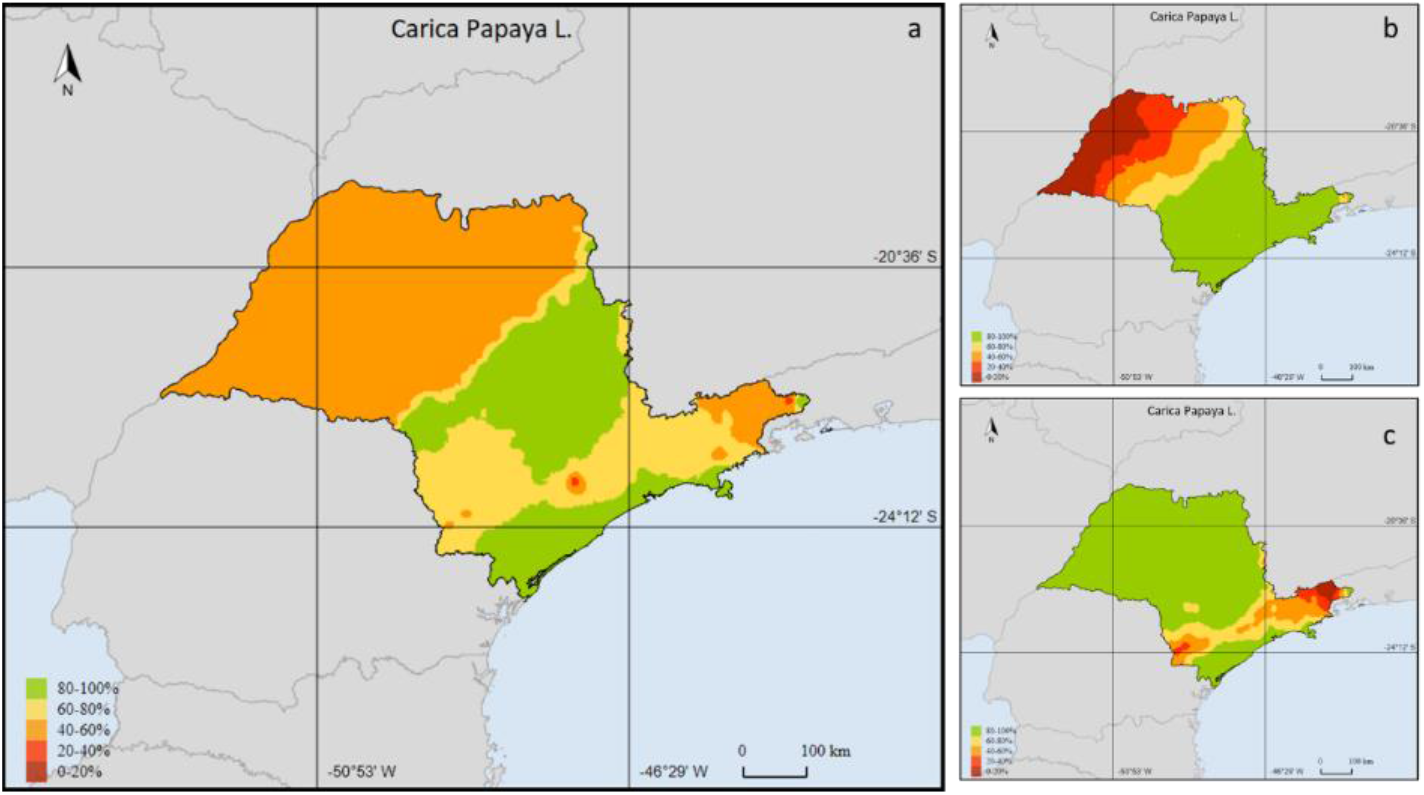
Suitable area in São Paulo State, Brazil, for Papaya cultivating using EtaHadgem ES future pessimist scenario. In (a): result with all variables, (b) annual water deficiency and (c) annual average temperature risk

**Table 8.**
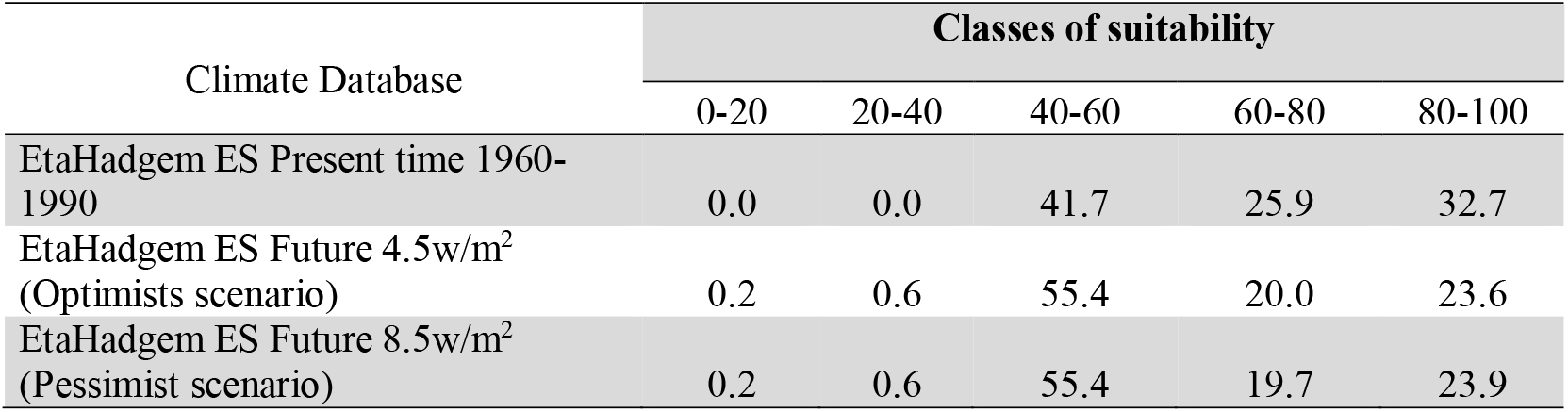
Surface of São Paulo are (in percentage) classified by level of suitability for Papaya production in present time and future climate change scenarios, using the climate regional model EtaHadgem ES

These results are consistent with Silva et al. (2013) and Carvalho et al. (2014) that specify the high water demands of the crop. The disequilibrium in water availability in the growth and production phase could affect negatively the papaya final production (Silva et al., 2013). Therefore, the high annual hydric deficiency projected by the scenarios studied here, mainly in the west of the State, will be an important limiting factor to restrict the papaya plantation.

Plant physiological stress due to water deficit may occur when precipitation is lower than the atmosphere evaporative demand (Yano et al., 2007), requiring more water to avoid declination in yield. In this context, one agronomic management technique that could be used in order to reduce the projected water deficiency is the irrigation. According to Coelho et al. (2011), irrigation is especially important during the papaya flowering phase because the water supply allows the complete process of flowering and fruiting, culminating in higher final production. Carvalho et al. (2014) discuss that with the advancement of technologies and the increasing demand for water, the search for more efficient irrigation methods has been accentuated. Therefore, drip irrigation, which is a practice of applying water through drops directly to the soil in the root zone, has emerged in the last two decades as a management that consume less resource and provide better results in productivity and quality.

### 3.5 Climatic Scientific Knowledge and BRAMAZOS software

The analyses using BRAMAZOS indicated decreases in suitable zone, mainly to Persimmon and Papaya cultivation, which could be mitigated by agronomic techniques as shading, agroforestry systems and irrigation. These management techniques, however, are not always accessible to smallholders because there are a lack access to technical information or financial support that could help them to make a more climate-resilient agriculture.

Nevertheless, the discussion of the potential results of the global climate change on agriculture is deeper than just plant/cultivate, or not, in a specific area. What is involved in this discussion, besides the food insecurity, is the vulnerability of a specific society group (in our case is the smallholders) which is exposed and is highly sensitive to changes in the climate pattern. The future (and actual) capacity of adaptive of the smallholders is normally low, generating serious social and economic challenges. Therefore, there is an urgent demand to create adaptation strategies and public politics to support smallholders activities, encouraging and providing access to the knowledge and scientific information, allowing current climate risk management and planning future activities to minimize the impact of the climate change linking adaptation and mitigation activities.

In this context, it is important the accessibility of climate information. To manage the climatic risk, it is important to understand current and future scenarios of climate and the relationship to the plant physiology, allowing decision-making over short and long time horizons, especially to non-scientific community. Even though the demand to scientific climatic information, it is remarkable that the information generated by the scientists are disconnected to the smallholder’s needs and understand (Donatti et al., 2017). The information more suitable for purpose and in a format that can be integrated in decisions, in an accessibility language, is crucial to help reduce risks and utilize this enormous potential. As discussed by Singh et al. (2018) there are few examples of long-term climate information being used in decisions at small and medium scale, and normally this information is inappropriate to decision-makers at the local scale, principally smallholders.

Therefore, BRAMAZOS is presented as an important tool to support efficient information to climate risk management, providing agroclimatic information that are capable of assist decision making, with the intention to reduce the climate impact on smallholders development and resources management issues.

### 3.6 Model Limitations

One important and delicate issue is the climate database. It is important to know the quality of the climate data as the climate series failures, since this data is the basis to run the model with satisfactory result. There are many series of climate data that are not consistent and present discontinuities, especially when we are using real data, collected by meteorological station. According to Ribeiro et al (2016) those inconsistences occurs because of the climate factors (as volcano, extreme precipitation for example) and collecting or recording data problems (issues in the process of measuring, processing, transfer, storing and transmitting the data). It can be caused by problems in the sensors that are measuring the meteorological element; computing and software questions; and, changes in the localization of station. All these factors may generate discontinuities and inhomogeneities in the climate time series that can result in misinterpretations of the analysed climate (Ribeiro et al. 2016).

Additionally, models that represent the climatic system and project future scenarios carry many uncertainties (Hawkins at al., 2009) which could also implies in uncertainties in the results of the future land availability to the crop. However, climate models projections are still the best tool to evaluate the impact of the climate change on several activities.

Although BRAMAZO software presents several functionalities, there is still a lack of knowledge that would be important to the smallholders, such as analyzing the best suitable areas for fruits quality (or agricultural products). This is especially important for crops that have altitude requirements. The quality of coffee beverage, for example, has a significant relationship with altitude (example).

We are still working on software development, and there are some issues that are not ready yet. Study the prevailing winds, for example, would be also desirable once we can indicate the orientation of the crop in cardinal points. Some orientation, for instance, has more probability to suffer frost. Finally, BRAMAZO do not included natural fluctuation on climate systems, such as El Nino and La Nina.

## 4 Conclusion

We presented here “Brazilian Mapping for Agricultural Zoning System” (BRAMAZOS) software which is a new system that was designed to support smallholders to manage their climatic risk on the production area, integrating climate (current and future), soil and plant physiology database. Samallholders are increasingly exposed to climate conditions and manage this risks are crucial to achieving agricultural production, reduce losses, increasefood security and promote sustainability. The information generated by BRAMAZOS presents a number of opportunities to investments in climate risk management, allowing adaptation option in a climate smart agriculture. Usually, the concept involved in climate risk management strategies includes activities designed in adaptation/mitigation; insurance (risk transfer) and risk confront (improving resilience of samallholders). All this approaches depends on the information and scientific knowledge transferred to the user. Therefore, the main idea of this new program is to transform scientific knowledge into useful information, indicating the potential impacts of climate indicators in the crops, allowing climate risk management, indicating the risk of crop failure and, thus, permitting smallholders to be better informed to initiate agriculture operations.

We used two temperate and one tropical fruit specie, which is commonly planted by smallholders, as an example to run the software in the current and future scenarios of climate change. The analyses using BRAMAZOS indicated that the temperate fruit analysed here, that require exposure to chilling conditions, could be more affected by temperatures increase, than the tropical specie analysed, that could be more affected by precipitation deficit. Persimmon that requires more winter chill than fig, presented more decrease in suitable zone. The reduction on the suitable zone could be mitigated by agronomic techniques as shading, agroforestry systems and irrigation.

## Acknowledgments

We acknowledge the financial support of The Brazilian National Council for Scientific and Technological Development (CNPQ), process n 4462-21/2015.

